# Epigenetics and island-mainland divergence in an insectivorous small mammal

**DOI:** 10.1101/2022.04.14.488253

**Authors:** Marie-Laurence Cossette, Donald T. Stewart, Amin Haghani, Joseph A. Zoller, Aaron B.A. Shafer, Steve Horvath

## Abstract

Geographically isolated populations, specifically island-mainland counterparts, tend to exhibit phenotypic variation in many species. The so-called island syndrome occurs when different environmental pressures lead to insular divergence from mainland populations. This phenomenon can be seen in an island population of Nova Scotia masked shrews (*Sorex cinereus*), which have developed a specialized feeding habit and digestive enzyme compared to their mainland counterparts. Epigenetic modifications, such as DNA methylation (DNAm), can impact phenotypes by altering gene expression without changing the DNA sequence. Here, we used a *de novo* masked shrew genome assembly and a mammalian methylation array profiling 37 thousand conserved CpGs to investigate morphological and DNA methylation patterns between island and mainland populations. Island shrews were morphologically and epigenetically different than their mainland counterparts, exhibiting a smaller body size. A gene ontology enrichment analyses of differentially methylated CpGs implicated developmental and digestive system related pathways. Based on our shrew epigenetic clock, island shrews might also be aging faster than their mainland counterparts. This study provides novel insight on phenotypic and epigenetic divergence in island-mainland mammal populations and suggests an underlying role of methylation in island-mainland divergence.

## Introduction

Insular populations tend to exhibit divergent traits compared to their continental counterparts (Adler & Levins, 1994; Foster, 1964; Goltsman et al., 2005; Meröndun et al., 2019; Novosolov et al., 2013; van Valen, 1973). This phenomenon is known as the island syndrome and refers to ecological and morphological differences between island and mainland populations (Baeckens & van Damme, 2020). Altered predation, competition, environment, and geographic isolation are thought to lead to phenotypic differences between mainland and insular populations (Baeckens & van Damme, 2020; Benítez-López et al., 2021; Lomolino et al., 2013).

Relatively little is known about the specific mechanisms causing phenotypic changes in island-mainland scenarios, whether it be genetic drift, natural selection, or plasticity (Baeckens & van Damme, 2020; Benítez-López et al., 2021). Miller et al. (2021) found heritable variations for deer mice for both mass and length associated with outlier loci between island-mainland populations and suggested that natural selection underlies such phenotypic divergence. Island-mainland populations, however, do not always exhibit clear signal of nuclear genetic divergence with divergent phenotypes (e.g., Meröndun et al., 2019; Vernaz, Malinsky, et al., 2021). In contrast, different epigenetic signatures, which include chemical modifications that regulate gene expression (Al Aboud et al., 2021), have been associated with substantial levels of divergence between species and populations exhibiting different phenotypes (Meröndun et al., 2019; Vernaz, Malinsky, et al., 2021) and unlike DNA, epigenetic modifications can respond quickly to environmental pressures (Anastasiadi et al., 2021). For example, differential methylation patterns associated with phenotypic traits have been noted for body size in sheep (*Ovis ammon*) (Cao et al., 2015) and lynx (*Lynx canadensis*) (Meröndun et al., 2019), for pigmentation levels in rabbits (*Oryctolagus cuniculus*) (Chen et al., 2021) and for breeding success in barn swallows (*Hirundo rustica*) (Saino et al., 2017).

Epigenetic modifications may take a variety of forms, these include DNA methylation, histone modifications and non-coding RNA activity (Moore et al., 2013), all of which can regulate gene expression without requiring changes in the DNA sequence (Razin & Riggs, 1980). Differential patterns of epigenetic modification have also been associated with early stages of speciation (Quadrana & Colot, 2016; Vernaz, Hudson, et al., 2021; Vernaz, Malinsky, et al., 2021; Vogt, 2017). DNA methylation (DNAm) involves the transfer of a methyl group to a cytosine base (Moore et al., 2013) and can occur throughout the genome but mostly occurs on CpG dinucleotide sites (Moore et al., 2013). Depending on where in the genome these modifications occur, they can either supress or activate gene expression (Dhar et al., 2021; Wan et al., 2015) and in some cases, methylation of only one CpG can inhibit the binding of transcription factors (Iguchi-Ariga & Schaffner, 1989; Watt & Molloy, 1988). CpG islands (CGI) are regions of the genome that contain a high frequency of CpGs relative to CpGs found in other regions of the mammalian genome (Illingworth & Bird, 2009). CGIs are also known to overlap most promoter regions (Gardiner-Garden & Frommer, 1987; Jones, 1999). These sites might be evolutionarily important in mammals since many CGIs are conserved between mice and humans (Deaton & Bird, 2011; Illingworth et al., 2010; Moore et al., 2013). It is theorized that over time differential methylation can lead to permanent heritable (cis) nuclear changes, referred to as genetic assimilation (Anastasiadi et al., 2021). This can happen when environmental conditions persist through generations favouring a specific phenotype, and over time resulting in selection for a permanent genetic basis (e.g., point mutations) for the phenotype previously produced from the plastic epigenetically-influenced phenotype (Anastasiadi et al., 2021).

In addition, DNAm levels, particularly hypermethylation, on specific CpG sites changes throughout an individual’s lifetime, making it possible to age individuals based on their methylation levels (Horvath, 2013; Lu et al., 2021; Prado et al., 2021; Wilkinson et al., 2021). These epigenetic *clocks* can predict chronological age in different mammalian species (Jasinska et al., 2021; Lu et al., 2021; Prado et al., 2021; Raj et al., 2021; Schachtschneider et al., 2021), and can be used to validate current aging techniques based on field observations (Robeck et al., 2021). Epigenetic clocks can also be used to predict demographic trends for conservation efforts (Prado et al., 2021) and have been shown to be elevated in inbred individuals, suggesting that inbreeding can lead to aging at a faster rate (Larison et al., 2021). Likewise, metabolic syndrome in humans (i.e., obesity) also results in increased epigenetic aging (Lee & Park, 2020; Nannini et al., 2019). It stands to reason that the differential selection pressures underlying island-mainland patterns might also alter the DNAm aging rate.

Shrews of the family Soricidae appeared about 30 to 40 million years ago (Churchfield, 1990). Hundreds of extant shrew species are now found on most continents (Churchfield, 1990; George, 1986, 1988) and have evolved unique traits such as echolocation (Chai et al., 2020; Forsman & Malmquist, 1988; Tomasi, 1979), venomous saliva (Kita et al., 2004) and the ability to dive underwater (Catania et al., 2008; Mendes-Soares & Rychlik, 2009). Shrews have the fastest mass-specific metabolic rate of all mammals (Ochocińska & Taylor, 2005) and are one of the smallest and shortest-lived mammals having a maximum lifespan of ∼17 months (Churchfield, 1990), making them important to study from an aging perspective (Lu et al., 2021). Shrews of the genus *Sorex* show differing trends in body size variation across their range; populations of some species at higher latitudes have smaller body sizes, which is counter to the predictions of Bergmann’s rule (Ochocińska & Taylor, 2003; Yom-Tov & Yom-Tov, 2005), whereas populations of other species are larger at more northerly latitudes (Huggins & Kennedy, 2018). *Sorex* shrews also exhibit morphological plasticity; the Eurasian common shrew (*Sorex araneus*), for example, manifests seasonal and reversible changes in skull morphology, body mass and organ size (Lázaro et al., 2017; Schaeffer et al., 2020; Taylor et al., 2013). Whether plastic or genetic in nature, these phenotypic variations throughout winter and latitude are generally explained as a response to limited resource availability (Lázaro et al., 2017; Ochocińska & Taylor, 2003; Schaeffer et al., 2020; Taylor et al., 2013; Yom-Tov & Yom-Tov, 2005).

Masked or Cinereus shrews (*Sorex cinereus*) are the most abundant shrew in North America (Nagorsen, 1996). The range of this species spans much of the northern United States and most of Canada (Nagorsen, 1996) with a varied diet consisting primarily of invertebrates (Bellocq et al., 1994; Churchfield, 1990; Whitaker John O. Jr. & French, 1984). One population of island dwelling masked shrews off the southern coast of Nova Scotia became isolated from the mainland during the late Wisconsin glaciation around 20,000 years ago (Roland, 1982; Stewart & Baker, 1992) and has developed a specialized feeding habit in response to their littoral environment (Stewart et al., 1989). Masked shrews on Bon Portage Island (BPI) are the only population of *Sorex* known to mainly feed on sand fleas (*Platorchestia platensis*) (MacPherson & Stewart, 2003; Stewart et al., 1989). *Platorchestia platensis* is a type of amphipod typically not consumed by insectivores, although Churchfield (1990) observed that the population of the lesser white-toothed shrew (*Crocidura suaveolens)* on St. Martin’s, Isles of Scilly, in the United Kingdom, also feed on amphipods. In the case of *S. cinereus*, electrophoresis assays by Stewart and Baker (1992) found that the BPI masked shrew population expressed a unique (slow) allele for Peptidase A and showed evidence of drift, but genetic differentiation across populations was low (Nei’s *D* ≌ 0.01). It was later hypothesized that due to the importance of sand fleas on BPI, the shrew’s digestive enzyme, Peptidase A, has undergone an adaptive response (MacPherson & Stewart, 2003). Here, using a *de novo* assembled and annotated genome, we built a masked shrew epigenetic clock and quantified differential methylation patterns between populations to examine the association between geographical isolation and epigenetic divergence. We hypothesized that island shrews will exhibit different phenotypes than mainland shrews based on the island syndrome and that divergent methylation patterns could underly these differences.

## Materials and methods

### Sampling, sexing, and aging shrews

Masked shrew samples were collected (Animal Care Certificate No. 26234) from five locations: Peterborough County (44°53′N, 78°05′W) in Ontario, Canada, and Sandy Cove (44°48′N, 66°08′W), North Mountain (45°13′N, 64°22′W), Long Island (44°19′N, 66°16′W) and Bon Portage Island (BPI) (43°28′N, 65°45′W) in Nova Scotia, Canada (Figure S1). All shrews were trapped between mid-July and late August to limit seasonal morphological differences, such as skull and body size (Lázaro et al., 2017, 2019, 2021). DNA was extracted from liver as this is where digestive enzymes are found; tail and fetal tissue were selected to help build the epigenetic clock (see Lu et al., 2021). Liver and tail samples produce high DNA yields and are easy to locate in shrews. We used a DNeasy Blood & Tissue Kit from QIAGEN. Shrews were left in dermestid beetle tanks for a week to remove all remaining tissue from the skulls. A Leica EZ4 microscope was used to assess age class for each shrew based on teeth wear according to the method of Pruitt (Pruitt, 1954; Rudd, 1955). Age in months was estimated based on the trapping date, the age class, and the known reproductive season of masked shrews assuming a maximum lifespan of ∼17 months (Churchfield, 1990). Shrews were sexed by gel electrophoresis of PCR amplicons of the Y chromosome-linked SRY gene (Cervantes et al., 2013; Matsubara et al., 2001) using custom primers (Table S1); sex assignments were subsequently validated with the methylation assay.

### Genome assembly and annotation

A single masked shrew from Peterborough County, Ontario had DNA and RNA extracted from the heart and liver immediately after euthanasia. We used the MagAttract HMW DNA Kit from QIAGEN and the Monarch Total RNA Miniprep Kit from New England BioLabs, respectively, and sent the extracts to The Centre for Applied Genomics, Toronto, Ontario, Canada, for sequencing. Short-read (SR) libraries were generated using Truseq PCR-free preparation and sequenced on two lanes on an Illumina HiSeq X platform with 150 bp paired-end reads. Linked reads (LR) were prepared using the 10X genome library after selecting fragments >20Kb using BluePippin. The linked reads were sequenced on one Illumina HiSeq X Lane. Twenty-five million RNA reads were sequenced on a HiSeq2500 with 126bp paired-end reads.

We used BBMap v. 35.8 (Bushnell, n.d.) to remove adapters and trim low-quality data from the fastq files. Contaminant screening was performed using Kraken2 v. 2.0.9-beta (Wood & Salzberg, 2014) and all unclassified sequences were kept. Kmergenie v. 1.7048 (Chikhi & Medvedev, 2014) was used to find the kmer size for downstream analysis. The genome was assembled using a tiered approach: 1) using only the SR data and Meraculous v.2.2.4 (Chapman et al., 2011) and 2) with only the LR data using Supernova v.2.1.1 (Weisenfeld et al., 2018) using the --maxreads filter set to “all”. The pseudohap2 style was selected for the assembly output. Following the assembly, both SR and LR assembly versions were merged via quickmerge v.0.3 (Chakraborty et al., 2016) with the LR as the backbone. Scaff10x v.5.0 (Ning et al., n.d.) was used to further polish the *S. cinereus* genome. Genome completeness was assessed using Benchmarking Universal Single-copy Orthologs (BUSCO) v.3.0.2 (Simão et al., 2015) by comparing the hybrid genome to highly conserved genes in mammals based on the *mammalia_odb9* dataset. The mitochondrial genome was assembled using MitoZ v.2.4-alpha (Meng et al., 2019). Genome annotation was conducted via GenSAS v.6.0 (Humann et al., 2019) using the assembled genome and RNAseq data. Here, repeat regions of the genome were identified using RepeatModeler v.1.0.1.1 (Smit & Hubley, n.d.-b) and RepeatMasker v.4.0.7 (Smit & Hubley, n.d.-a). Masked shrew liver and heart RNA reads were mapped to the genome using HISAT2 v.2.1.0 (Kim et al., 2019). The resulting BAM file was used by BRAKER v.2.1.1 (Hoff et al., 2019) and EVidenceModeler v1.1.1 (Haas et al., 2008) to predict features in the genome. Functional annotation to find common protein sequences between the masked shrew and the common shrew, *Sorex araneus* (GenBank assembly accession: GCA_000181275.2), and other vertebrate mammals was done using BLAST v.2.11.0 (Altschup et al., 1990), SwissProt v.2.11.0 (Bairoch & Apweiler, 1998) and InterProScan v.5.44-79.0 (Jones et al., 2014) databases.

### Epigenetic clocks

DNA methylation profiles for each sample (n = 48, Table S2) were assayed on the HorvathMammalMethylChip40, a novel Infinium array that profiles up to 37,000 highly conserved CpGs across the genome (Arneson et al., 2022). The SeSaMe normalization method was used to define beta values for each probe (Zhou et al., 2018). Unsupervised hierarchical clustering based on inter-array correlation coefficients revealed that samples cluster by tissue type. The methylation array measurements can assess the sex of each sample based on the X chromosomal CpGs. One fetal sample was removed due to inaccurate sexing. Only probes that aligned to the masked shrew genome (bwa-mem) were retained for downstream analysis (n = 29,609). Elastic net regression models (alpha = 0.5, 10-fold internal cross-validation) were applied to the DNAm data (covariates) and the chronological age estimate (dependent variable). The resulting multivariate regression model will be referred to informally as the masked shrew epigenetic clock. The alpha value of the glmnet R software package (Friedman et al., 2010) was set to 0.5 to be midpoint between a ridge and lasso type regression. Elastic net regressions help prevent overfitting the model to the data by selecting variables (CpGs) and shrinking them simultaneously (Zou & Hastie, 2005). Three different models were used as per Lu et al. (2021); 1) no transformation to the chronological age, 2) log-transformed chronological age (log(x+1)), and 3) square root transformed chronological age (sqrt(x+1)) with offsets of 1 year to avoid negative values for our fetal samples. To assess the accuracy of our epigenetic clock models we used a leave-one-out (LOO) cross-validation method. This method consists of training the epigenetic clock models with a subset of the data and testing how well it can predict other observed data. We report the LOO estimates of the Pearson correlation between age and its DNAm based estimates as well as the median absolute error (MAE) between DNAm age and observed age (estimated chronological age in years).

### Statistical analyses and functional gene enrichment

We ran a series of models in R v.1.3.959 (R Core Team, 2021). Using linear models, we separately regressed body size, weight, and skull length against location, age and sex (i.e., phenotype ∼ location + age + sex) of 33 masked shrews (Table S2). Here, location was a categorical variable of island or mainland (see Table 1). Additionally, we extracted the residuals from the epigenetic clock for each sample and ran a Welch Two Sample T-test using location and sex to test if different groupings epigenetically aged at comparable rates. Positive residuals were indicative of an epigenetically older individual than the linear model would predict.

**Table 1.**
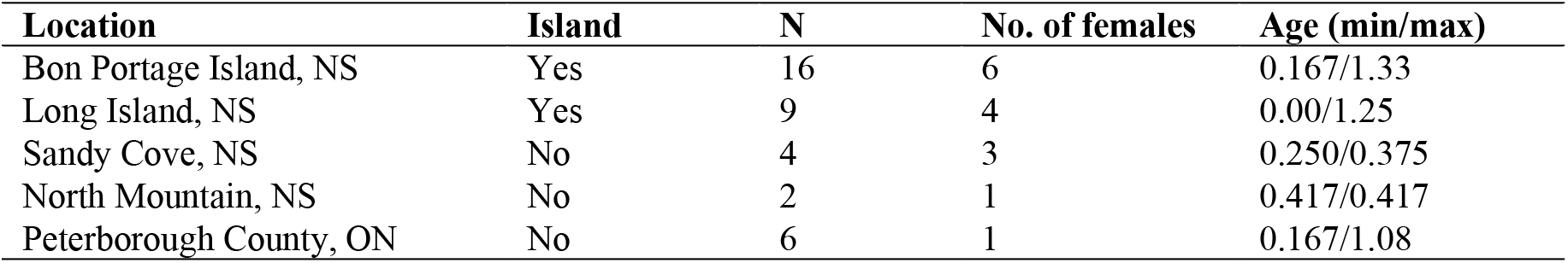
Trapping location of samples, N = total number of samples, number of females, minimum and maximum age. Age is represented as the estimated age in months divided by 12. Note, age of 0.00 represents fetus.

Epigenome-wide association studies (EWAS) were conducted using the limma package v.3.44 (Ritchie et al., 2015) with 44 methylomes (Table S2); note we removed the fetal samples from this analysis to avoid biased results associated with developmental genes and pathways. Using the β values of aligned probes, we ran the following model: DNAm ∼ location + sex + Age + tissue including animal ID as a random effect. This model was run two times; once with BPI and Long Island grouped together as ‘Island’ in the model with all other locations assigned as ‘Mainland’ and once with BPI versus all other populations. The latter grouping was meant to isolate possible unique dietary adaptations of BPI shrews. The island versus mainland model was also run with only Long Island samples to test for a potential bias of having more BPI samples in the island grouping.

A *p* value < 2e-6 was set according to the Bonferroni-corrected threshold as the cutoff for significance to be conservative (Johnson et al., 2010). The location of probes in relation to the gene annotations were identified using the ChIPseeker (Yu et al., 2015) and GenomicFeatures (Lawrence et al., 2013) packages. Enriched pathways associated with the significant CpGs for the EWAS models were identified with rGREAT v.1.20 (McLean et al., 2010) using the human Hg19 genome annotation to allow for comparisons (Jasinska et al., 2021; Prado et al., 2021; Schachtschneider et al., 2021). We accounted for bias resulting from the design of the mammalian array by using 14,290 CpGs that mapped to our shrew gene annotations as background. Following Schachtschneider et al. (2021), from these probes we filtered and kept the top 500 hypermethylated and top 500 hypomethylated sites based on *p* value to use for the enrichment analysis.

## Results

### Genome assembly and annotations

We assessed the quality and completeness of our genome assembly to ensure confidence in our downstream analyses. The SR and LR assembly statistics are provided in Table S3; the final hybrid genome was 2.66 Gb with an N50 of 1.99 Mb and L50 of 291 scaffolds. The GC content was 42.88% with over 86% of the genome being in scaffolds over 50 Kb and the largest scaffold being around 22 million base pairs. Close to 95% of BUSCOs were identified in this assembly (Table S3): 3504 of 4101 were complete single-copies, 74 were complete duplicates and 301 were fragmented. Repeats represented 38.60% of the genome (Table S4) with long interspersed nuclear elements (LINEs) and unclassified repeats representing 20.17% and 15.39% of the genome respectively. The functional annotation pipeline identified 20,067 protein coding genes (Table S3). The metrics assessed were comparable to other genome assemblies using similar sequencing approaches (e.g., Etherington et al., 2020; Wolf et al., 2022). The raw sequence data have been deposited in the Short Read Archive (SRA) under accession number PRJNA826195. The assembly has been deposited at DDBJ/ENA/GenBank (accession no. pending).

### Epigenetic clocks

We built a shrew methylation clock to assess epigenetic aging rates between different shrew populations (n = 24 from islands versus n = 12 from mainland). The methylation data consisted of 33 tail, 11 liver, and 3 fetal tissue DNA samples selected from 36 different individuals (Table S2). One fetal sample was dropped due to quality control metrics (wrong sex). The average age of our samples was seven months; the youngest shrews being fetuses and the oldest one being an estimated 16 months (Table 1). Out of the 37,000 probes, 29,609 were aligned to our masked shrew genome. We constructed and cross-validated epigenetic clocks for different masked shrew tissue types, which included liver, tail, fetus, and pan-tissue clock. The final clocks were based on the methylation profiles of 26 CpGs, similar to clocks from other species such as elephants or cats (e.g., Prado et al., 2021; Raj et al., 2021), our clocks were highly accurate with a LOO cross validation estimate of the Pearson correlation between age and its predicted value r >= 0.91 and a median absolute error of MAE = 0.13 years for the multi-tissue clock, representing approximately a 7-week difference between the estimated DNAm age and the teeth-based chronological age. The liver DNAm clock had the lowest MAE of 0.07 years (∼3 weeks), followed by the multi-tissue and tail clocks with MAE around 0.1 (∼5 weeks) (Figure 1).

**Figure 1.**
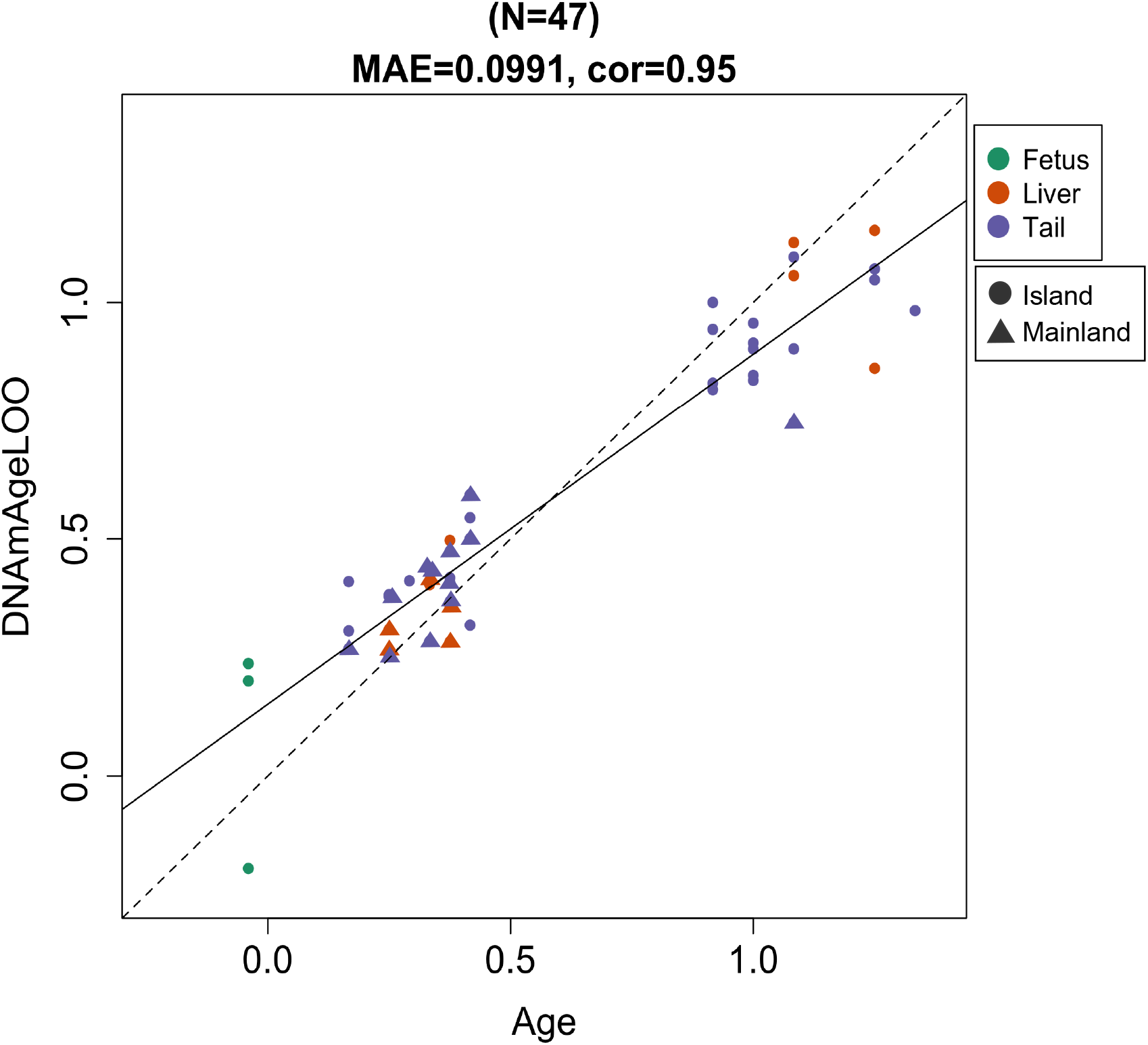
Cross-validation study of pan-tissue epigenetic clock for masked shrews. Leave-one-sample-out (LOO) estimate (y-axis, in units of years) versus chronological age (x-axis, in unit of years). The linear regression of epigenetic age is indicated by a solid line while the diagonal line (y = x) is depicted by a dashed line. “MAE” represents the mean absolute error and “cor” represents the correlation coefficient

### Island-mainland differences

Morphological and epigenetic age differences between 21 island and 12 mainland masked shrew populations (Table S2) were examined using linear regression. Mass, body length and skull length differed between the mainland and island samples (Figure 2). Our models including age, sex, and island-mainland covariates explained a high percentage of variance: skull length (Adj. R^2^ = 0.79), weight (Adj. R^2^ = 0.53) and body length (Adj. R^2^ = 0.49; Table 2). In all models, location was the variable that correlated most to the change in body size and weight (*p* value < 0.01; Table 2). All size metrics were negatively correlated to being on an island (Table 2, Figure 2). Age was most often positively correlated to body size whereas sex did not have an effect (Table 2).

**Table 2.** Summary statistics for the linear regression models for weight, body length and skull length of mainland versus island masked shrews. Sex values are relative to females and location relative to mainland populations. The β-coefficient (estimate), 95% confidence interval (CI), p-value and adjusted R^2^ are provided.

|  | Estimate | 95% CI | P value | Adj. R<sup>2</sup> |
| --- | --- | --- | --- | --- |
| Model 1: Weight ~ Age + Sex + Location |  |  |  | 0.53 |
| Age | 1.15 | 0.21 to 2.08 | 0.018 |  |
| Sex | 0.01 | -0.61 to 0.64 | 0.962 |  |
| Location | -2.09 | -2.79 to -1.39 | <0.001 |  |
| Model 2: Body length ~ Age + Sex + Location |  |  |  | 0.49 |
| Age | 8.43 | 3.36 to 13.50 | 0.002 |  |
| Sex | -0.20 | -3.61 to 3.20 | 0.903 |  |
| Location | -10.35 | -14.15 to -6.54 | <0.001 |  |
| Model 3: Skull length ~ Age + Sex + Location |  |  |  | 0.79 |
| Age | -0.31 | -0.93 to 0.32 | 0.321 |  |
| Sex | -0.21 | -0.63 to 0.21 | 0.319 |  |
| Location | -1.92 | -2.39 to -1.45 | <0.001 |  |

**Figure 2.**
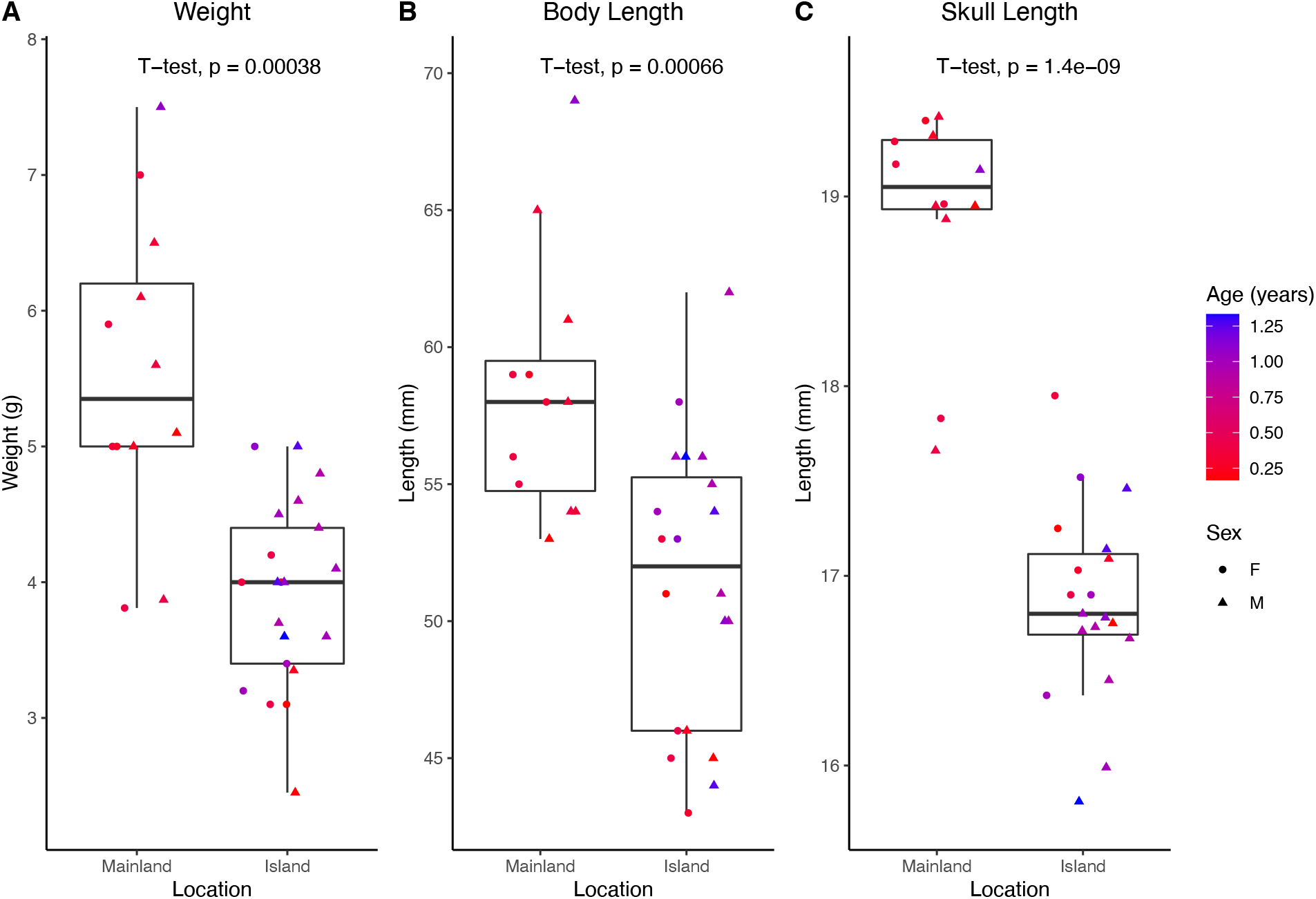
Boxplot of **A** weight (g), **B** body length (mm) and **C** skull length (mm) between mainland and island masked shrew populations.

Epigenetic clock residuals were slightly elevated on islands indicative of individuals epigenetically older than predicted (t = −1.95, df = 39.73, *p* value = 0.06); residual variation was higher on the island (mean of 0.01) compared to the mainland (mean of −0.02). There was no difference in residuals among the sexes (t = −0.97, df = 40.2, *p* value = 0.34) or age groups (yearlings versus older) (t = −0.21, df = 11.79, *p* value = 0.84).

### EWAS Island effects

EWAS were conducted using the methylation data consisting of the 33 tail, 11 liver samples (Table S2) to identify differentially methylated genes and pathways between island and mainland masked shrews. A total of 1067 differentially methylated CpG sites (482 hypermethylated and 585 hypomethylated relative to mainland) were identified (*p* value < 2e-6) between our island and mainland populations (Figure 3A). The most divergent CpG was on the *EWS* exon (*p* value = 3.35e-21) a gene that encodes a multifunctional protein involved in various cellular processes such as gene expression, cell signaling, and RNA processing and transport. Top differentially methylated CpGs were located on or near genes associated to developmental and adipose-related processes: *MEST* exon (*p* value = 3.34e-14), *SIM1* intergene (*p* value = 3.79e-13), *PHIP* intergene (*p* value = 1.03e-12) and metabolism: *ZBTB7C* promotor (*p* value = 8.45e-14) and *ATPA* exon (*p* value = 6.37e-16) (Table 3, Figure 3A, 3B). The top hypermethylated CpGs were associated to three different fibroblast growth factor receptors pathways (*p* value < 5e-16) (Figure 3C). Other significantly enriched pathways included synaptic and adipose-related processes (Figure 3C).

**Table 3.** Top 5 genes near outlier CpGs for island versus mainland EWAS and BPI versus all other populations. Functions of each gene are abbreviated from Uniprot.

| Gene name | CpG | Direction | Location (p) | Putative Function |
| --- | --- | --- | --- | --- |
| Island-mainland |  |  |  |  |
| EWS exon | cg11703808 | pos | 3.35e-21 | Transcriptional repressor and might play a role in the tumorigenic process. |
| PO4F3 intron | cg00152260 | pos | 1.23e-19 | Transcriptional activator. Involved in the auditory system development. |
| UBP54 intron | cg22063749 | neg | 3.27e-19 | Has no peptidase activity. |
| TRI47 exon | cg25115601 | neg | 3.12e-16 | Mediates the ubiquitination and proteasomal degradation of CYLD. |
| ATPA exon | cg24102071 | neg | 6.37e-16 | Produces ATP from ADP |
| BPI |  |  |  |  |
| SMO exon | cg20625795 | neg | 1.03e-30 | Associates with the patched protein (PTCH) to transduce the hedgehog's proteins signal. |
| ZO1 exon | cg03673751 | neg | 2.00e-21 | Related to movement of substances through the paracellular space. Plays a role in the regulation of cell migration. |
| PRP8 promoter | cg25476805 | neg | 7.60e-19 | Plays role in pre-mRNA splicing. |
| CBX5 intergenic | cg23036219 | neg | 1.12e-18 | Associated with epigenetic repression. Involved in the formation of functional kinetochore. |
| ZBT20 intergenic | cg00903722 | neg | 3.32e-17 | Transcription factor that may be involved in hematopoiesis, oncogenesis, and immune responses. Plays a role in postnatal myogenesis. |

**Figure 3.**
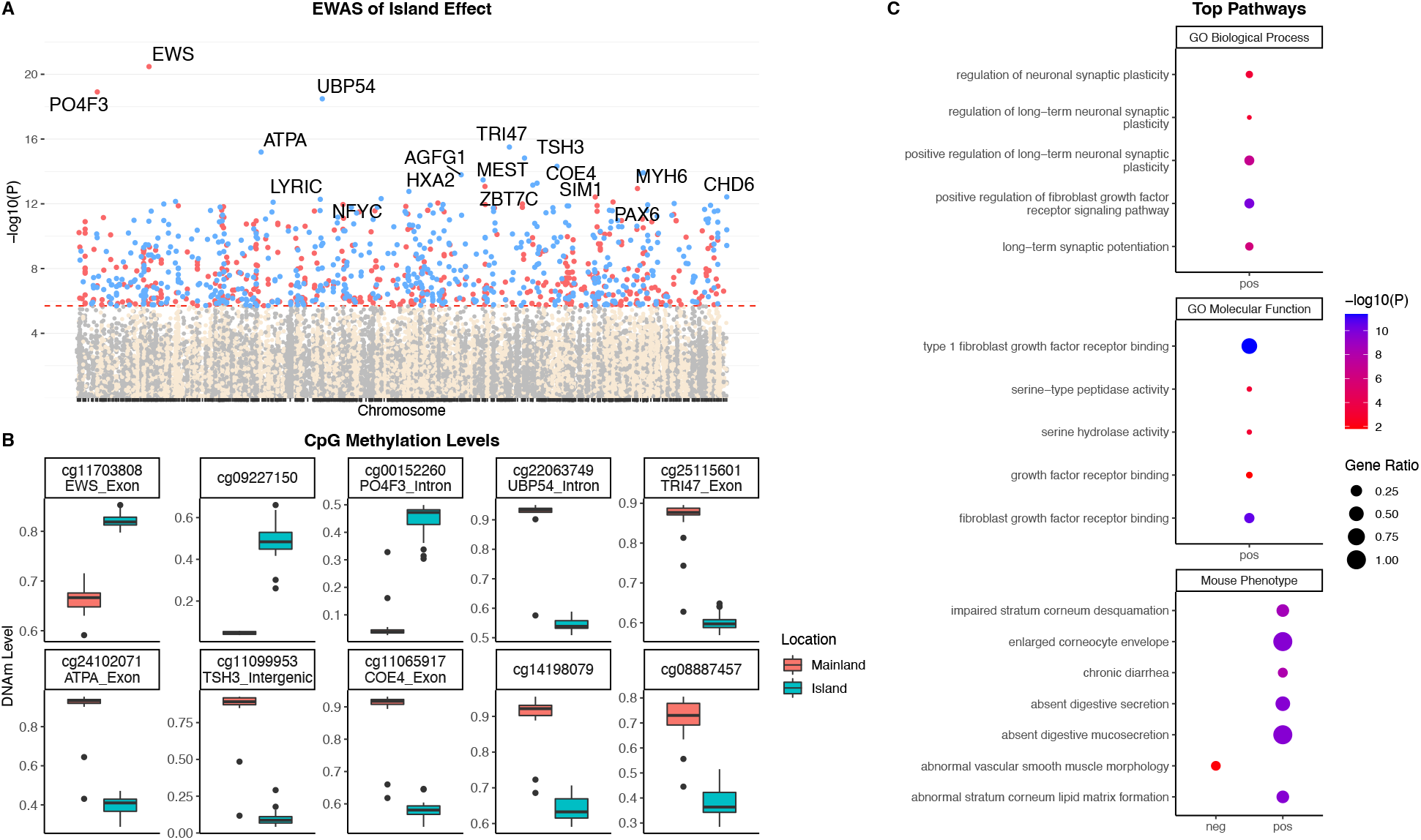
Epigenome-wide association (EWAS) of the multivariate regression model (location, age, sex and tissue) for masked shrew. **A** Manhattan plots of the EWAS for location (island versus mainland). The coordinates are estimated based on the alignment of 29,609 Mammalian array probes to our masked shrew genome assembly. The direction of associations with < 2e-6 (red dotted line) is highlighted by red (hypermethylated) and blue (hypomethylated) colors relative to mainland methylation levels. Top 15 CpGs are indicated by their neighboring genes. **B** DNAm levels of island samples versus mainland for the top 10 significant CpGs (*p* value) associated to location. **C** Enrichment analysis of the top CpGs with positive (hypermethylated) and negative (hypomethylated) correlations to island populations. The gene-level enrichment analysis was carried out using the GREAT software. Background probes were limited to 14,290 probes that had shrew gene annotations. The top ontologies with most significance pathways for island samples were selected based on Bonferroni corrected p-values.

Isolating BPI identified 485 significant CpGs (*p* value < 2e-6), 308 were hypomethylated and 177 were hypermethylated compared to other populations (Figure 4A). The top significant hypomethylated CpG in BPI samples was on the *SMO* exon (*p* value = 1.42e-30; Table 3, Figure 4A, 4B). Top CpGs were most often hypomethylated in BPI island samples (Table 3, Figure 4A, 4B). Pathway analysis highlighted that differentially methylated sites in BPI shrews compared to other populations seem to affect digestive and intestinal phenotypes and functions (Figure 4C). Additionally, other phenotypes related to skull morphology, more specifically the mandible, teeth, and ear canal showed pathway enrichment in BPI shrews (Fig 4C). When BPI was removed from the island-mainland model (i.e. only Long Island), digestive enzymes were not identified as being a significantly enriched pathway (Figure S2).

**Figure 4.**
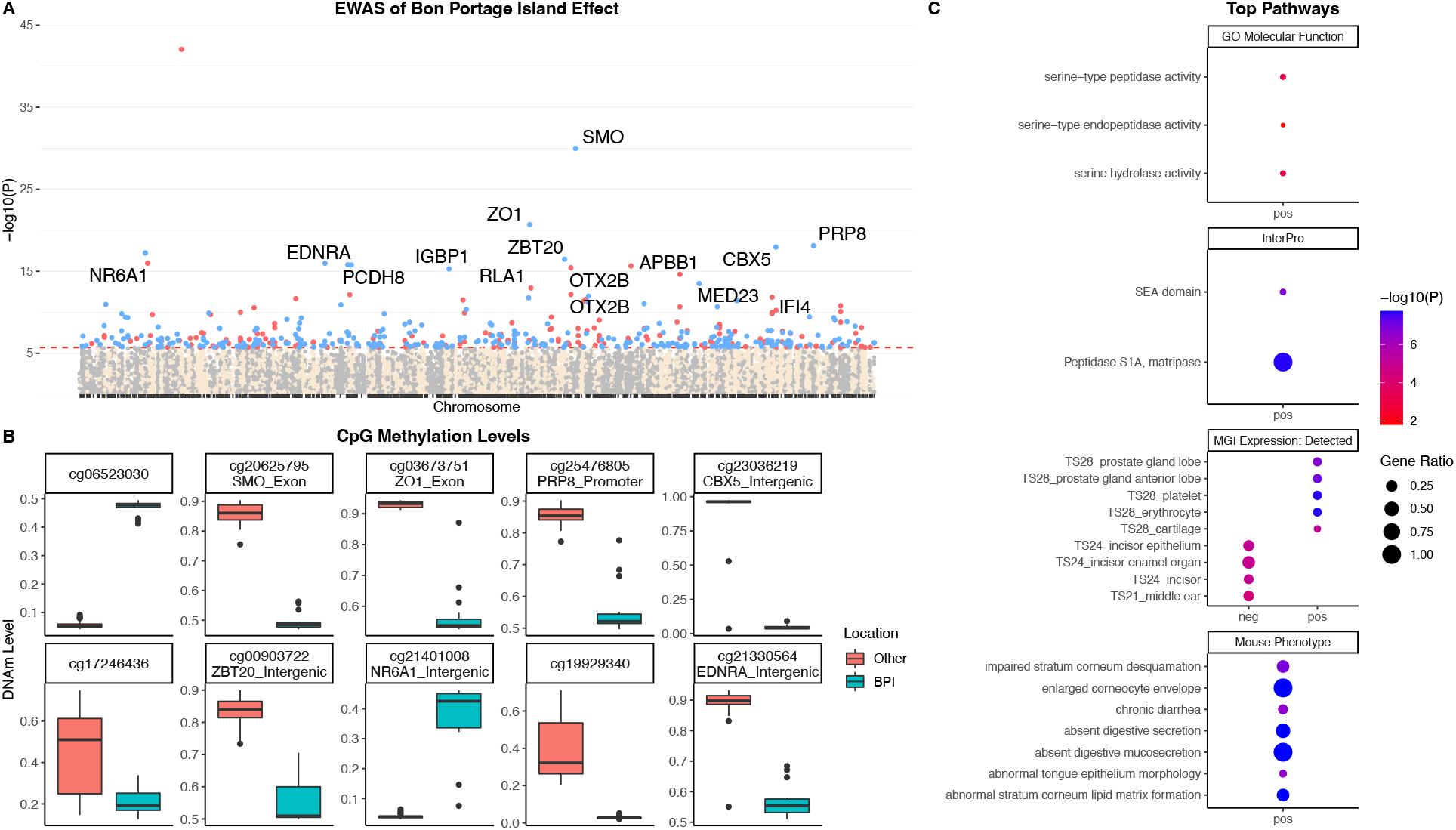
Epigenome-wide association (EWAS) of the multivariate regression model (location, age, sex and tissue) for masked shrew. **A** Manhattan plots of the EWAS for location (Bon Portage Island (BPI) versus others). The coordinates are estimated based on the alignment of 29,609 Mammalian array probes to our masked shrew genome assembly. The direction of associations with < 2e-6 (red dotted line) is highlighted by red (hypermethylated) and blue (hypomethylated) colors relative to non-BPI methylation levels. Top 15 CpGs are indicated by their neighboring genes. **B** DNAm levels of BPI samples versus others for the top 10 significant CpGs (*p* value) associated to location. **C** Enrichment analysis of the top CpGs with positive (hypermethylated) and negative (hypomethylated) correlations to BPI. The gene-level enrichment analysis was carried out using the GREAT software. Background probes were limited to 14,290 probes that had shrew gene annotations. The top 4 ontologies with most significance pathways for BPI were selected based on Bonferroni corrected p-values.

## Discussion

The island syndrome describes the distinctive morphological differences between island-mainland populations due to different ecological and environmental pressures (Baeckens & van Damme, 2020). We examined whether phenotypic divergence was associated with epigenetic modifications, specifically DNAm, which alter gene expression without changing the DNA sequence (Razin & Riggs, 1980). Various authors have argued that methylated cytosines reduce gene expression in promotors by causing chromatin structures to condense which influences the binding of transcription factors (Moore et al., 2013; Pappalardo & Barra, 2021). However, methylation in other parts of the genome, e.g., intragenic, or distal regions, can both repress and induce gene expression (Anastasiadi et al., 2018). Our analysis showed that differential methylated patterns are associated with distinct phenotypes between island and mainland masked shrew populations. Over 29,000 CpG methylation probes aligned to our genome, consistent with other mammalian studies (Larison et al., 2021; Pinho et al., 2021; Schachtschneider et al., 2021; Sugrue et al., 2021), and a robust epigenetic clock was generated (Figure 1). The differential methylation between island-mainland populations showed a remarkable overlap between enriched pathways and measured phenotypic differences. While some association assessments suffer from false positives and “story-telling” (Pavlidis et al., 2012), there was an unambiguous connection to the measured body phenotypes (Figure 2), dietary differences (MacPherson & Stewart, 2003; McAlpine, 2009; Stewart et al., 1989), and the observed enriched pathways (Figure 3C, 4C). Thus, the clear morphological and dietary island-mainland differences appear to be, in part, under epigenetic influence.

The island syndrome describes the distinctive morphological differences between island-mainland populations due to different ecological and environmental pressures (Baeckens & van Damme, 2020). We examined both phenotypic divergence and epigenetic modifications in masked shrews, specifically DNAm which has the potential to increase phenotypic plasticity within an individual (Vogt, 2017). Highly diversified DNAm methylation among populations can be a strategy to take advantage of unpredictable environments (Anastasiadi et al., 2021); moreover, epigenetic modifications have been shown to allow isogenic lines to adjust their phenotype to maximize fitness (van Egeren et al., 2018). Bon Portage Island masked shrew populations have lower genetic diversity than their mainland counterparts (Stewart & Baker, 1992), but likely have a broader ecological niche due to reduced interspecific competition; masked shrews on mainland Nova Scotia are broadly sympatric with six other species of shrews, and diet studies demonstrate that these soricids differentially exploit available invertebrate food resources (Whitaker & French, 1984). Without standing genetic variation, differential methylation among offspring could be a bet-hedging strategy to allow for exploration and exploitation of a range of food resources on the island (Anastasiadi et al., 2021). Further, DNAm can theoretically lead to genetic modifications through the process of genetic assimilation where methylated cytosines have a higher frequency of mutagenesis resulting in novel alleles (Anastasiadi et al., 2021). Differential methylation is an effective strategy that individuals in genetically impoverished populations may use to generate phenotypic diversity. Thus, we hypothesize that BPI shrews are in the process of methylation-induced genetic assimilation, with the high frequency of the unique peptidase A allele in this population being one outcome of this process.

### Phenotypic and epigenetic divergence

DNA methylation regulates gene expression thereby contributing to phenotypic plasticity (al Aboud et al., 2021; Kooke et al., 2015; Razin & Riggs, 1980), which can lead to phenotypic variation among individuals subject to different environmental conditions (Anastasiadi et al., 2021; Kooke et al., 2015; Meröndun et al., 2019). Here, insular masked shrews exhibited different phenotypes than their mainland counterparts; morphologically, this presented as decreased body length, mass, and skull length on islands. Body size is a heritable polygenic trait in mammals (Posbergh & Huson, 2021; Postma, 2014) with many underlying mechanisms, often making it difficult to clearly understand the role of each gene or pathway involved (e.g., Anderson et al., 2022). Cytosine methylation levels measured in highly conserved stretches of DNA have been linked to mammalian traits such as maximum lifespan (Haghani et al., 2021) and has been suggested as a universal mechanism to regulates body mass and morphological development in animals in other association studies (Cao et al., 2015; Haghani et al., 2021). Genetic variation in quantitative trait loci (QTL) almost certainly play a role in phenotypic variations between insular and mainland populations – and might also influence methylation (J. T. Bell et al., 2011; Smith et al., 2014). We suggest that pooled genome sequencing approaches (e.g., island versus mainland samples) are a feasible option to test for nuclear genome associations of this nature.

Differential methylated pathways between island and mainland shrew populations for all tissue types included fibroblast growth factors (FGF), which are crucial for animal body development (Oulion et al., 2012), and adipose-related genes and processes. Haghani *et al*. (2021) also found that across mammals, species weight was correlated to development and adipose-related pathways, possibly due to DNAm affecting the expression of adipose tissue and regulation of lipid storage (Ma & Kang, 2019). In masked shrews, CpGs on the homolog to mesoderm-specific transcript protein (*MEST*) exon and single-minded homolog 1 (*SIM1*) intergenic region were hypo and hypermethylated respectively in island shrews compared to mainland shrews. *MEST* is known to be correlated to adipocytes in humans (Karbiener et al., 2015) and mice (Takahashi et al., 2005), while *SIM1* is associated with food intake regulation and obesity (Holder et al., 2004; Yang et al., 2006), both possibly affecting body mass in shrews. Differential methylation of genes such as *SIM1* might affect shrews’ metabolism and ultimately body size by regulating the expression of hormones involved in signaling for food intake. Further, island shrews had a hypomethylated CpG in the pleckstrin homology domain-interacting protein (*PHIP*) intergenic region, a probable regulator of insulin-like growth factor signaling pathways (Farhang-Fallah et al., 2002), which reinforces the idea that insulin growth factors are involved in mammalian life-history variation (Swanson & Dantzer, 2014). The overall associations between DNAm and genes related to body composition, possibly indicates that methylation inhibits or activates various genes during developmental stages in shrews, but future studies are needed to better understand such mechanisms.

Masked shrew insular populations appear to have decreased body mass compared to their mainland counterparts; BPI masked shrews occur in high densities (Downie, 1986; Stewart et al., 1989; Telfer, 1984), which should lead to small animals having larger body size to better compete for resources (Juette et al., 2020). However, the discrepancy with this expected phenotype for island shrews might be due to their diet. Previous studies have shown that food scarcity contributes to the selection of smaller body size in Palearctic shrews, which might also be the case for Nearctic populations (Ochocińska & Taylor, 2003; Yom-Tov & Yom-Tov, 2005). Churchfield (2002) found that in locations where higher biomass resources are scarce, shrews tend be smaller in size and survive by consuming less nutritional but more abundant sources of food. This might be the case for BPI shrews feeding on sand fleas. Supporting this idea, top EWAS genes involved in energy homeostasis and metabolism were differentially methylated between mainland-island populations (Table 3). For example, ATP Synthase F1 Subunit Alpha (*ATPA*) catalyzes the synthesis of ATP, the main molecule used to store and transfer energy in cells (Schapira, 2006). Similarly, the zinc finger and BTB domain–containing 7c (*ZBTB7C)* gene promotor was also differentially methylated in island shrews. *ZBTB7C* is an important metabolic regulating gene for blood-glucose homeostasis during fasting (Choi et al., 2019).

BPI shrews also showed differentially methylated sites associated with various processes related to the digestive system, including peptidase activity. This is of significance as previous studies noted that masked shrews on BPI have developed specialized littoral feeding habits (MacPherson & Stewart, 2003; Stewart et al., 1989). Previous work showed that BPI shrews possess a unique polymorphism for the digestive enzyme Peptidase A (Stewart & Baker, 1992), which potentially allows shrews to easily digest the sand fleas’ exoskeleton (McAlpine, 2009). Digestive system related pathways being in the top enrichment category indicate that these dietary habits seem to be one of the most significant molecular differences between this insular population and others and support earlier allozyme work (MacPherson & Stewart, 2003; McAlpine, 2009; Stewart & Baker, 1992). Our work remains a correlative, but future genomic assays can be used to test our genetic assimilation hypothesis, while mouse models or common-garden experiments could be used to assess the localization of these CpGs and phenotypic associations to provide information on possible mechanism of enriched regions on masked shrew methylomes.

### Epigenetic age

DNA methylation and chronological age can be used to build epigenetic clocks to identify genes associated with aging and predict how methylation patterns change throughout an individual’s lifetime (Larison et al., 2021; Lu et al., 2021; Prado et al., 2021; Wilkinson et al., 2021). Epigenetic clocks have been constructed for a wide range of species, showing consistent trends with CpGs associated to developmental genes and pathways (Lu et al., 2021; Pinho et al., 2021; Raj et al., 2021; Schachtschneider et al., 2021). Second generation clocks, such as the DNAm GrimAge, account for variables such as chronological age, health factors and mortality, to predict lifespan and healthspan (C. G. Bell et al., 2019; Lu et al., 2019). In humans, DNAm GrimAge has been studied in relation to various comorbidities and social factors (Lu et al., 2019; McCrory et al., 2021). Inbreeding has been associated with decreasing longevity and an increase in health concerns in various species (Keller & Waller, 2002; Skotarczak et al., 2020; Yordy et al., 2020) and recently was linked to accelerated DNAm aging in zebras (*Equus quagga*) (Larison et al., 2021). We used these patterns to explore accelerated aging as a component of island-mainland divergence.

Masked shrews tend to live on average 14 months; accurately aging small mammals such as shrews can be difficult since they are such small creatures and current methods rely on somewhat subjective classification based on teeth wear, body length measurements and weight (Pruitt, 1954; Rudd, 1955). Our epigenetic clock validated the current aging techniques used in the field, and overall, the clock was highly predictive (R = 0.95; MAE = 0.10) (Figure 1). Further, when we examined the clock model residuals, island shrews appeared to age slightly faster than mainland shrews. Parasites can affect hosts’ life-history traits through the energetic investment of maintaining an immune system response (Cooper et al., 2012; Cowan et al., 2009; Morand & Harvey, 2000), possibly resulting in faster biological aging. Group size has been shown to be correlated with prevalence and intensity of parasitism (Coté & Poulin, 1995). The population density of shrews on BPI is extremely high, at least in some years (Downie, 1986; Stewart et al., 1989; Telfer, 1984) possibly resulting in an increased incidence of various parasites and pathogens. Overall, this study reports divergent phenotypic trends between mainland-island masked shrew populations, and more broadly support the role of epigenetics in shaping phenotypic divergence. Future research should investigate epigenetic aging in insular populations, especially when inbreeding (Larison et al., 2021) or phenotypic divergence (i.e., this study) is present.

## Acknowledgments

We want to thank the reviewers for the valuable insights and comments provided. This work was supported by CanSeq150 (CGEn) (ABAS, DTS and MLC); Ontario Graduate Scholarship (MLC), Natural Sciences and Engineering Research Council of Canada (NSERC) Alexander Graham Bell Canada Graduate Scholarship-Master’s (MLC) and NSERC Discovery grants (ABAS and DTS); Compute Canada Resources for Research Groups (ABAS); Canadian Foundation for Innovation: John R. Evans Leaders Fund (ABAS); Ontario Early Researcher Awards (ABAS). SH was supported by the Paul G. Allen Frontiers Group. We wish to acknowledge that the work for this study was carried out on the traditional territory of the Mississauga (Michi Saagiig) Anishnaabeg and the Mi’kmaq People. We are grateful to have had the opportunity to work on this land and thank the First Peoples for their care, stewardship, and teaching.

## Data accessibility statement

Raw sequence reads are deposited in the SRA: PRJNA826195

Genome is deposited in NCBI: Pending

Scripts and normalized β values for each probe and sample uploaded on GitLab: https://gitlab.com/WiDGeT_TrentU/graduate_theses/-/tree/master/Cossette.

## Benefit-sharing statement

Benefits Generated: Benefits from this research accrue from the sharing of our data and results on public databases as described above.

## Author contribution

**Marie-Laurence Cossette:** Methodology, Software, Formal analysis, Investigation, Data curation, Writing – original draft preparation, Visualization, Project administration, Funding acquisition. **Don T. Stewart:** Conceptualization, Methodology, Resources, Writing – review and editing, Supervision, Funding acquisition. **Amin Haghani:** Writing – review and editing, Software, Data curation. **Joseph Zoller:** Writing – review and editing, Software, Data curation. **Aaron B.A. Shafer:** Conceptualization, Methodology, Validation, Investigation, Resources, Writing – original draft preparation, Supervision, Project administration, Funding acquisition. **Steve Horvath:** Conceptualization, Methodology, Software, Validation, Resources, Data curation, Writing – review and editing, Supervision, Project administration, Funding acquisition.

