## Supplemental for "Epigenetics and island-mainland divergence in an insectivorous small mammal"

MOLECULAR ECOLOGY

### Table of Contents:


| | |
|---|---|
| **Table S1** | Page 1 |
| **Table S2** | Page 1-2 |
| **Table S3** | Page 2 |
| **Table S4** | Page 3 |
| **Figure S1** | Page 3 |
| **Figure S2** | Page 4 |
| **Figure S3** | Page 5 |


**Table S1**. Master mix formula. Custom primer: 5′-CATGGTGTGGGCTCGCAATC-3 and 5′-CTGCCTGTAGTCTCTGTGCC-3'. PCR cycles as follow: denature at 93°C for 2 minutes, run 30 cycles of 15 seconds at 93°C, 20 seconds at 60°C and 20 seconds at 72°C.

| **Reagents** | **Conc. of stock solution** | **Volume per individual reaction** |
|---|---|---|
| Buffer | 5X | 4 µL |
| dNTPs | 2mM | 2 µL |
| $MgCl_2$ | 50mM | 1 µL |
| BSA | 3mg/mL | 1 µL |
| Primer | 10µM | 0.8 µL |
| *Taq* polymerase | 5U/µL | 0.2 µL |
| DNA | | 2 µL |
| Water | | 11 µL |
| Total volume (before DNA) | | 20 µL |

**Table S2.** Breakdown of sample information. n = represents the total number of samples used for each analysis. *represents second sample from a same individual.

| **SampleID** | **Tissue** | **Sex** | **Teeth class** | **Age (months)** | **Location** | **Used for epigenetic clock (n = 47)** | **Used for body size analysis (n = 33)** | **Used for EWAS (n = 44)** |
|---|---|---|---|---|---|---|---|---|
| 1213 | Tail | F | 2 | 3.5 | Bon Portage Island, NS | yes | yes | yes |
| 1215 | Tail | M | 1 | 2 | Bon Portage Island, NS | yes | yes | yes |
| 1216 | Tail | M | 2 | 3 | Bon Portage Island, NS | yes | yes | yes |
| 1217 | Tail | F | 2 | 5 | Bon Portage Island, NS | yes | yes | yes |
| 1222 | Tail | M | 3 | 11 | Bon Portage Island, NS | yes | yes | yes |
| 1223 | Tail | F | 1 | 5 | Bon Portage Island, NS | yes | yes | yes |
| 1225 | Tail | F | 1 | 2 | Bon Portage Island, NS | yes | yes | yes |

| | | | | | | | | |
|---|---|---|---|---|---|---|---|---|
| 1226 | Tail | M | 3 | 12 | Bon Portage Island, NS | yes | yes | yes |
| 1227 | Tail | M | 3 | 12 | Bon Portage Island, NS | yes | yes | yes |
| 1228 | Tail | M | 3 | 12 | Bon Portage Island, NS | yes | yes | yes |
| 1229 | Tail | F | 3 | 12 | Bon Portage Island, NS | yes | yes | yes |
| 1231 | Tail | M | 3 | 11 | Bon Portage Island, NS | yes | yes | yes |
| 1232 | Tail | F | 3 | 12 | Bon Portage Island, NS | yes | yes | yes |
| 1233 | Tail | M | 3 | 11 | Bon Portage Island, NS | yes | yes | yes |
| 1234 | Tail | M | 4 | 16 | Bon Portage Island, NS | yes | yes | yes |
| 1235 | Tail | M | 3 | 11 | Bon Portage Island, NS | yes | yes | yes |
| 1335 | Tail | M | 2 | 5 | North Mountain, NS | yes | yes | yes |
| 1336 | Tail | F | 2 | 5 | North Mountain, NS | yes | yes | yes |
| SC1 | Tail | M | 2 | 4.5 | Peterborough County, ON | yes | no | yes |
| SC2 | Tail | M | 1 | 2 | Peterborough County, ON | yes | yes | yes |
| SC3 | Tail | M | 4 | 13 | Peterborough County, ON | yes | yes | yes |
| SC4 | Tail | M | 2 | 4 | Peterborough County, ON | yes | yes | yes |
| SC5 | Tail | M | 2 | 4 | Peterborough County, ON | yes | no | yes |
| SC6 | Tail | F | 2 | 4.5 | Peterborough County, ON | yes | yes | yes |
| SC001 | Tail | F | 2 | 4.5 | Sandy Cove, NS | yes | no | yes |
| SC002 | Tail | F | 2 | 3 | Sandy Cove, NS | yes | no | yes |
| SC003 | Tail | F | 2 | 4 | Sandy Cove, NS | yes | no | yes |
| SC004 | Tail | M | 2 | 3 | Sandy Cove, NS | yes | no | yes |
| LI001 | Tail | M | 4 | 15 | Long Island, NS | yes | no | yes |
| LI002 | Tail | F | 4 | 13 | Long Island, NS | yes | no | yes |
| LI003 | Tail | F | 2 | 4.5 | Long Island, NS | yes | no | yes |
| LI004 | Tail | M | 4 | 13 | Long Island, NS | yes | no | yes |
| LI005 | Tail | M | 4 | 15 | Long Island, NS | yes | no | yes |
| SC1 | Liver* | M | 2 | 4.5 | Peterborough County, ON | yes | yes | yes |
| SC5 | Liver* | M | 2 | 4 | Peterborough County, ON | yes | yes | yes |
| SC001 | Liver* | F | 2 | 4.5 | Sandy Cove, NS | yes | yes | yes |
| SC002 | Liver* | F | 2 | 3 | Sandy Cove, NS | yes | yes | yes |
| SC003 | Liver* | F | 2 | 4 | Sandy Cove, NS | yes | yes | yes |
| SC004 | Liver* | M | 2 | 3 | Sandy Cove, NS | yes | yes | yes |
| LI001 | Liver* | M | 4 | 15 | Long Island, NS | yes | yes | yes |
| LI002 | Liver* | F | 4 | 13 | Long Island, NS | yes | yes | yes |
| LI003 | Liver* | F | 2 | 4.5 | Long Island, NS | yes | yes | yes |
| LI004 | Liver* | M | 4 | 13 | Long Island, NS | yes | yes | yes |
| LI005 | Liver* | M | 4 | 15 | Long Island, NS | yes | yes | yes |
| LI002_FET1 | Fetus | F | 0 | 0 | Long Island, NS | yes | no | no |
| LI002_FET2 | Fetus | M | 0 | 0 | Long Island, NS | yes | no | no |
| LI002_FET3 | Fetus | F | 0 | 0 | Long Island, NS | no | no | no |
| LI002_FET4 | Fetus | M | 0 | 0 | Long Island, NS | yes | no | no |

**Table S3.** Assembly statistics for masked shrew genome assemblies.

| | **Short-read assembly** | **Linked-read assembly** | **Hybrid assembly** |
|---|---|---|---|
| **Assembly size** | 1.92 Gb | 2.66 Gb | 2.66 Gb |
| **N50/L50** | 2.77 Kb / 175,854 scaffolds | 1.99 Mb / 291 scaffolds | 1.99 Mb / 291 scaffolds |
| **N90/L90** | 1.23 Kb / 607,012 scaffolds | 0.16 Mb / 1,755 scaffolds | 0.16 Mb / 1,753 scaffolds |
| **Number of scaffolds** | 779,274 | 84,561 | 84,558 |
| **Largest scaffold** | 91,699 | 22,233,027 | 22,233,027 |
| **% Genome in scaffolds > 50 Kb (%)** | 0.03 | 86.15 | 86.14 |
| **GC content (%)** | 42.63 | 42.88 | 42.88 |
| **BUSCOs identified** | | | 3,879 (95%) |
| **Number of protein coding genes** | | | 20,067 |

MOLECULAR ECOLOGY

**Table S4.** Summary of repeats in the masked shrew genome.

| | Length (bp) | Percentage of genome (%) |
|---|---|---|
| **SINE** | 53,159,155 | 2.10 |
| **LINE** | 509,850,416 | 20.17 |
| **LTR elements** | 2,632,939 | 0.10 |
| **DNA elements** | 665,221 | 0.03 |
| **Unclassified** | 389,117,414 | 15.39 |
| **Small RNA** | 0 | 0 |
| **Satellites** | 0 | 0 |
| **Simple repeats** | 20,505,599 | 0.81 |
| **Low complexity** | 0 | 0 |
| **Total** | 975,930,744 | 38.60 |

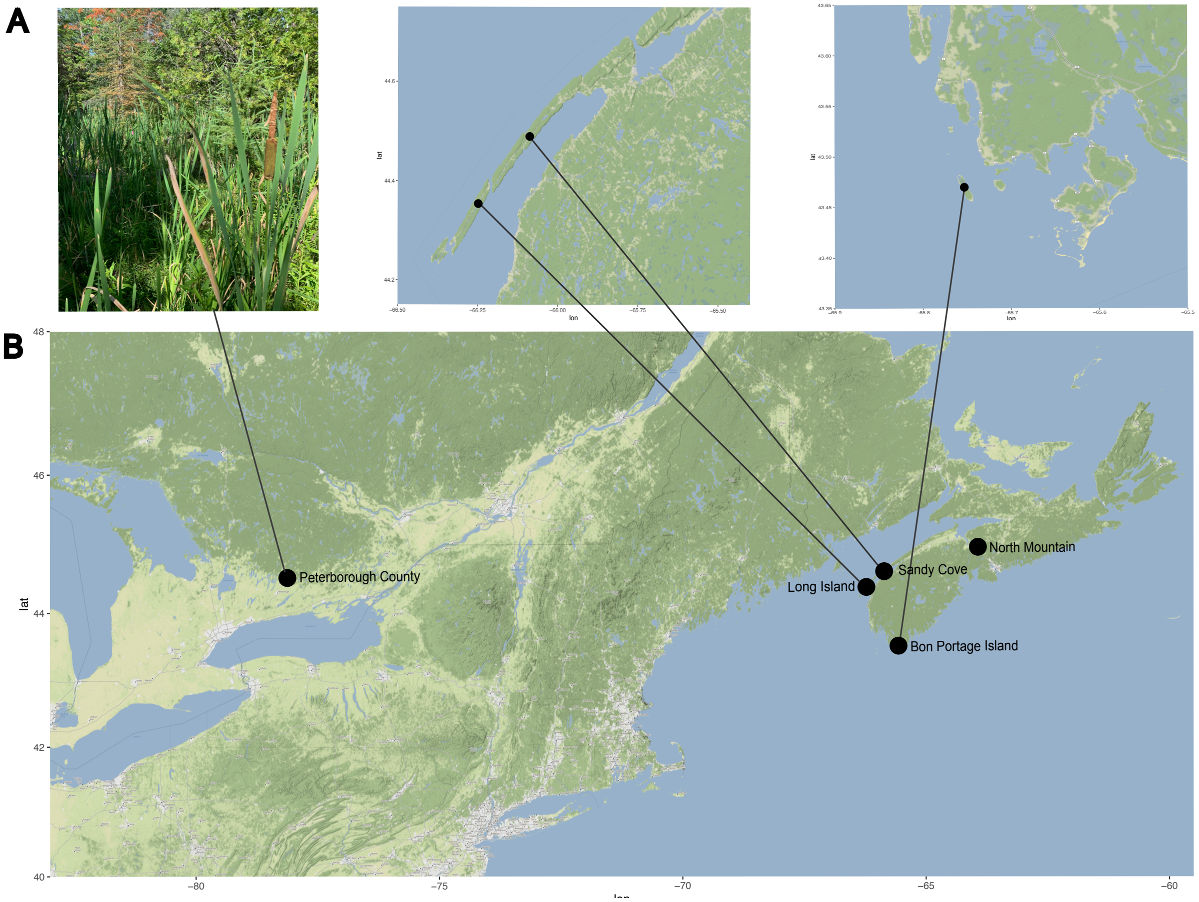


**Figure S1.** Trapping information for our masked shrew samples. **A** Example picture of the type of ecosystem where all traps were set. **B** Map of trapping locations in Ontario and Nova Scotia. Traps in Peterborough, Sandy Cove and North Mountain were set on the mainland. Bon Portage is located ~ 2.5 km from the mainland and Long Island is separated by ~ 750 m of water from the mainland. Both islands require a ferry or boat to reach.

MOLECULAR ECOLOGY

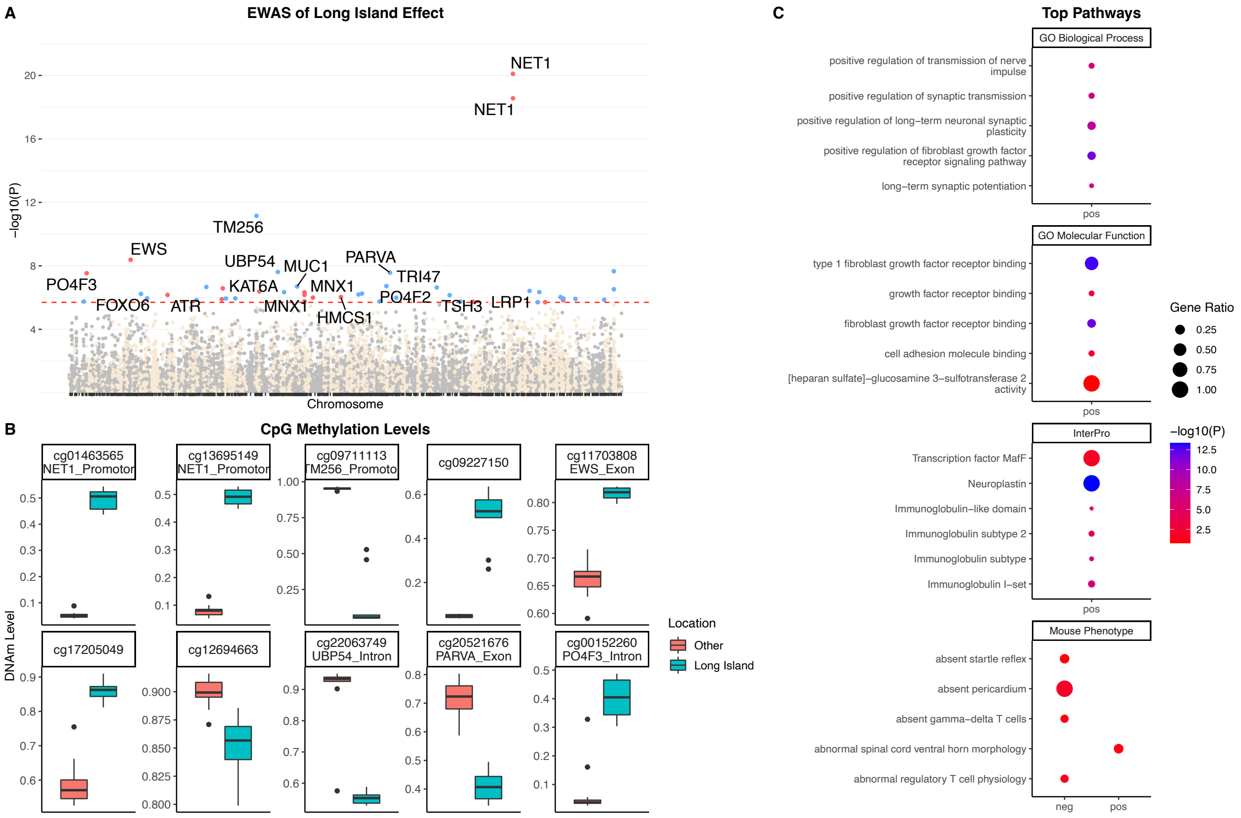


**Figure S2.** Epigenome-wide association (EWAS) of the multivariate regression model (location, age, sex and tissue) for masked shrew. **A** Manhattan plots of the EWAS for location (Long Island vs others). The coordinates are estimated based on the alignment of Mammalian array probes to our masked shrew genome assembly. The direction of associations with < 2e-6 (red dotted line) is highlighted by red (hypermethylated) and blue (hypomethylated) colors relative to non-Long Island methylation levels. Top 15 CpGs are indicated by their neighboring genes. **B** DNAm levels of Long Island samples versus others for the top 10 significant CpGs (p value) associated to location. **C** Enrichment analysis of the top CpGs with positive (hypermethylated) and negative (hypomethylated) correlations to Long Island. The gene-level enrichment analysis was carried out using the GREAT software. Background probes were limited to 14290 probes that had shrew gene annotations. The top 4 ontologies with most significance pathways for Long Island were selected based on Bonferroni corrected p-values.

MOLECULAR ECOLOGY

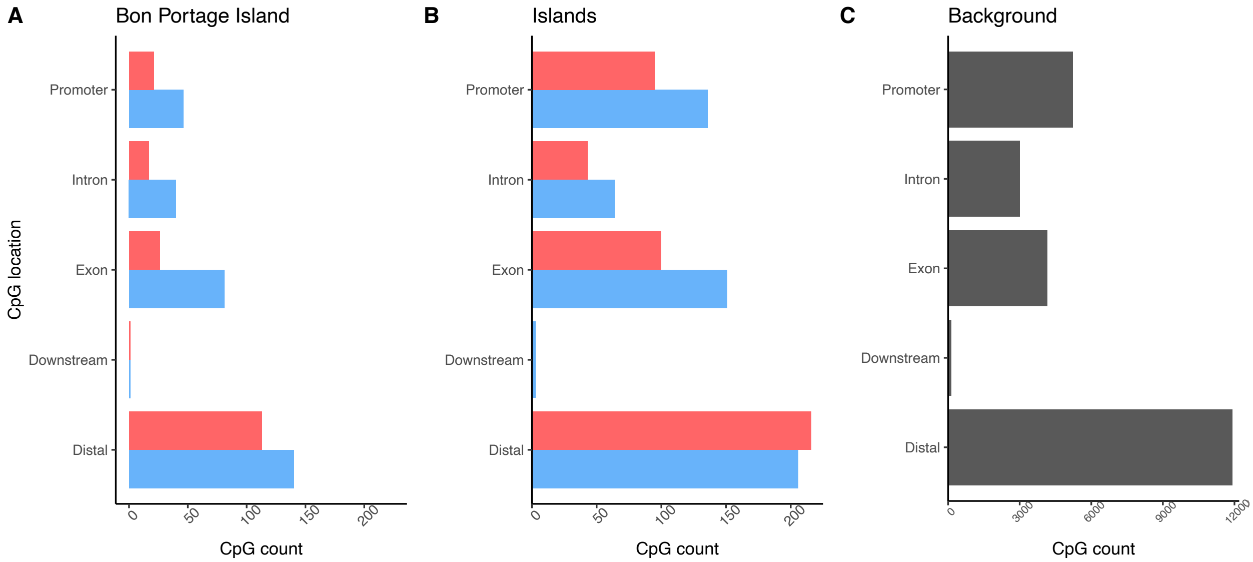


**Figure S3.** Location of probes aligned to the masked shrew genome for, **A** Bon Portage Island shrews relative other populations, **B** Island shrews relative to mainland shrews and **C** all probes. Blue bars represent hypomethylated CpGs and red bars represent hypermethylated CpGs.
